# Aging under immunosuppression reshapes human immune compartments and lowers clinical alloreactivity after heart transplantation

**DOI:** 10.64898/2026.02.25.681707

**Authors:** Kaushik Amancherla, Phillip Lin, B. Lakshitha A. Perera, Nelson Chow, Quanhu Sheng, Hasan K. Siddiqi, Eric Farber-Eger, Quinn S. Wells, Jane E. Freedman, Kelly H. Schlendorf, Ravi Shah, Eric R. Gamazon

## Abstract

Solid-organ transplantation in aging recipients represents a unique opportunity to study how age-related immunity in the context of non-specific immunosuppression strategies balances infection, malignancy, and rejection. Heart transplantation is an exemplar platform, as routine endomyocardial biopsy for rejection surveillance is the clinical “gold standard” regardless of clinical status. Here, we undertook the largest granular study to date to characterize the association between increasing recipient age at heart transplantation with acute allograft rejection and age-related cell-specific transcriptomic changes in circulating immune cells. This single-center retrospective cohort study evaluated individuals undergoing heart transplantation between July 2013 and December 2023 at Vanderbilt University Medical Center. Eligible participants were aged ≥18 years. A subset of individuals underwent single-cell RNA-sequencing of circulating immune cells. Among 799 adults, each one standard deviation increase in recipient age was associated with a ∼17% lower odds of allograft rejection (adjusted OR 0.83, 95% CI 0.71-0.98). In 40 individuals who underwent single-cell RNA-sequencing of circulating immune cells, increasing recipient age was associated with increases in *CD4+* and *CD8+* memory T cell subsets, monocytes, and NK cells. Furthermore, genes upregulated with increasing recipient age were associated with enrichment for pathways involved in immunosenescence and chronic low-grade inflammation while downregulated genes suggested decreased protein synthesis. These findings have clinical implications for an aging transplant population and support a more personalized approach to immunosuppression.

## INTRODUCTION

Age-related changes in immune function can substantially influence clinical disease response and progression, including infection, malignancy, and cardiovascular disease^1-3^. While immune dysregulation has been recognized in oncology and informs precision therapeutic approaches (via patient risk stratification, predicting immune-related adverse events, etc.)^4-8^, the situation in organ transplantation is complex, where age-related decline in immune function may be beneficial in inducing allograft tolerance^9^. Heart transplantation (HT) offers a unique opportunity to delineate alloimmune responses associated with aging, as routine endomyocardial biopsy for rejection surveillance is the “gold standard” for diagnosing allograft rejection^10-12^. Management of immunosuppression in an aging recipient population must balance allograft rejection, infection, and malignancy^10,13^, and heterogeneity across individuals may drive alloimmune responses central to these clinical outcomes^9^. Nevertheless, studies aimed at understanding age-related immune function during immunosuppression at the cellular level remain limited and are critical to clarifying the balance between immune tolerance and over-immunosuppression. Here, we sought to characterize the relationship between recipient age and risk of acute cardiac allograft rejection within the first-year post-HT.

## RESULTS

Our study overview is outlined in **Figure 1**. Of 799 adult recipients (30% women), 31% experienced allograft rejection within 1-year post-HT (the highest risk period), spanning different immune mechanisms (cellular, antibody-mediated, and mixed rejection). Full recipient and donor demographics are outlined in **Table 1**. In multivariable regression, each one SD increase in recipient age was associated with a ∼17% lower odds of allograft rejection (adjusted OR 0.83, 95% CI 0.71-0.98; **Figure 2**) after adjusting for donor age, recipient sex, and induction immunosuppression therapy. In a sensitivity analysis including only HT recipients who survived at least 90 days post-HT (N=768), each 1-SD increase in recipient age remained significantly associated with lower odds of allograft rejection (adjusted OR 0.81, 95% CI 0.69-0.95).

**Figure 1.**
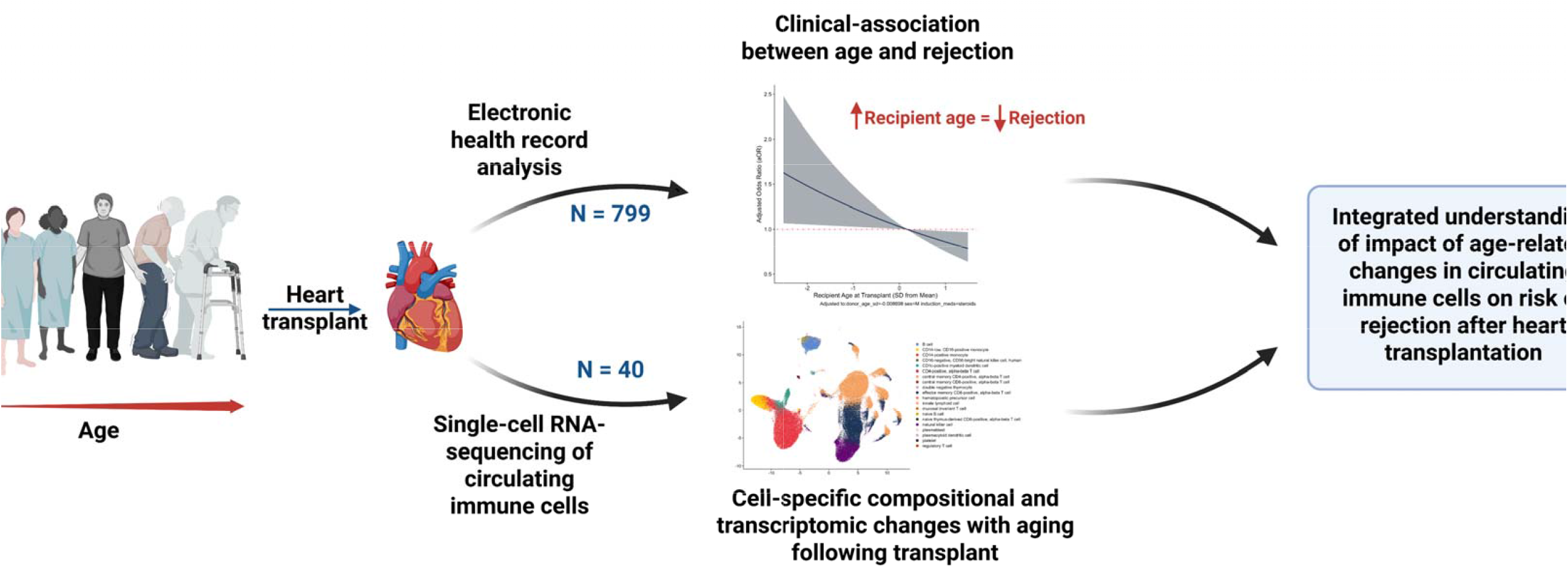
Study overview.

**Table 1.** Clinical characteristics of heart transplant donors and recipients.

| **Characteristic** | **N = 799<sup>1</sup>** |
| --- | --- |
| **Recipient age at transplant (years)** | 55 (45, 63) |
| **Recipient gender** |  |
| Female | 237 (30%) |
| Male | 562 (70%) |
| **Allograft rejection (within 1-year)** | 247 (31%) |
| **Rejection type** |  |
| Acute cellular rejection | 196 (25%) |
| Antibody-mediated rejection | 45 (5.6%) |
| Mixed rejection | 6 (0.8%) |
| No rejection | 552 (69%) |
| **Induction medications** |  |
| Basiliximab | 275 (34%) |
| Steroid-only | 423 (53%) |
| Thymoglobulin | 101 (13%) |
| **Donor age (years)** | 31 (24, 38) |
| **Donor cause of death** |  |
| Donation after brain death | 646 (81%) |
| Donation after circulatory death | 153 (19%) |

<sup>1</sup>Median (IQR); n (%)

**Figure 2.**
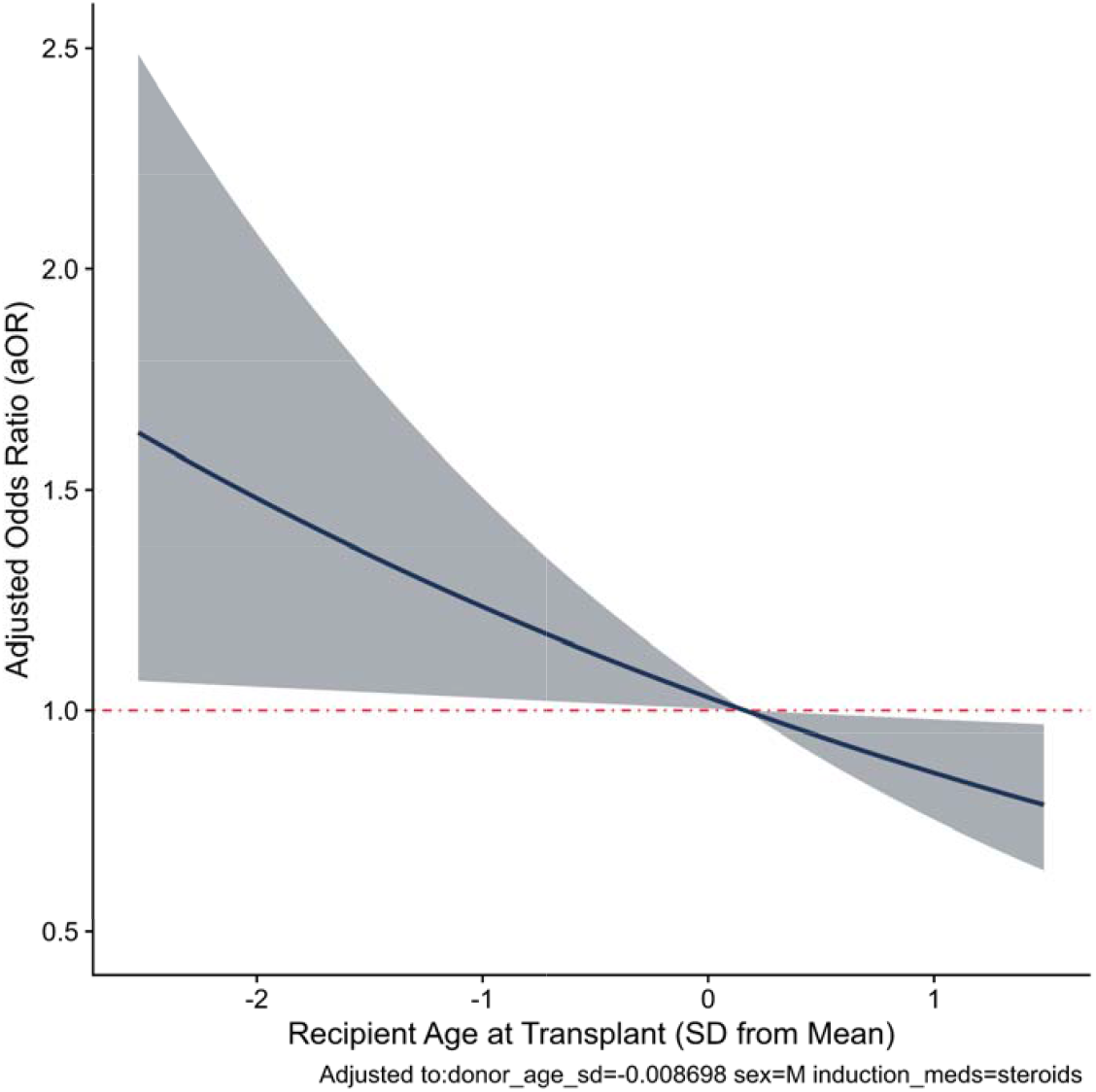
Association between recipient age (represented in standard deviations) and acute rejection within 1-year following heart transplant, assuming a linear relationship with the 95% CI highlighted around the adjusted odds ratio (aOR). The red dashed line represents an aOR of 1.

To identify alterations in immune mechanisms that occur with aging in immunosuppressed individuals, we performed single-cell RNA sequencing in HT recipients (recently reported by our group^14^) spanning 19-73 years of age (median age 53 years, 45% women; see **Supplemental Tables** for full clinical demographics). Among peripheral blood mononuclear cells (PBMCs) from 40 HT recipients, 152,106 cells spanning 20 unique clusters passed quality control (**Figure 3A**). With increasing recipient age, we identified 963 differentially abundant neighborhoods (FDR < 0.05; **Figure 3B, Supplemental Tables**), including significant increases in *CD4*+ and *CD8+* memory subsets, *CD14+* and *CD16+* monocytes, and NK cells.

**Figure 3.**
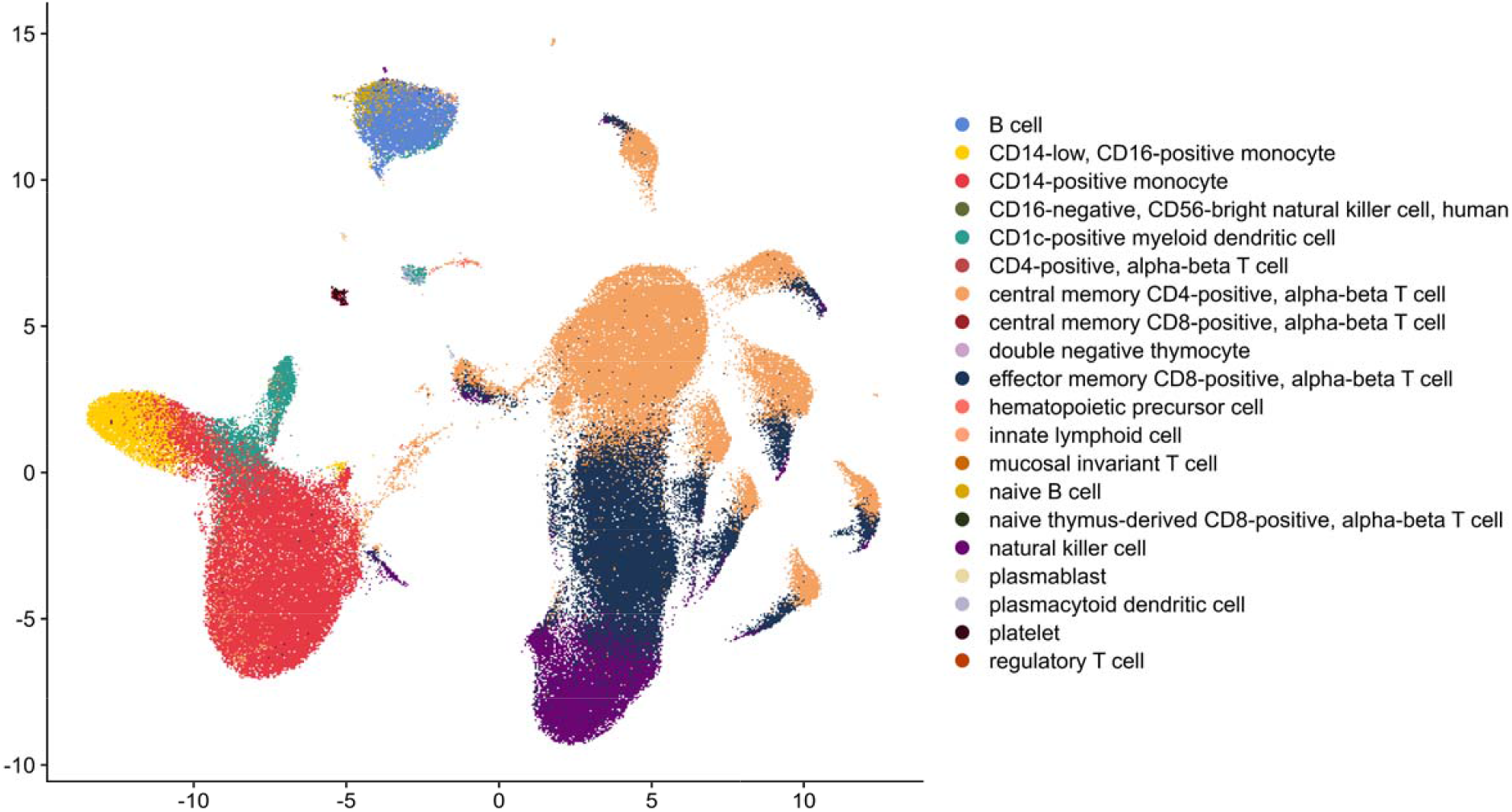
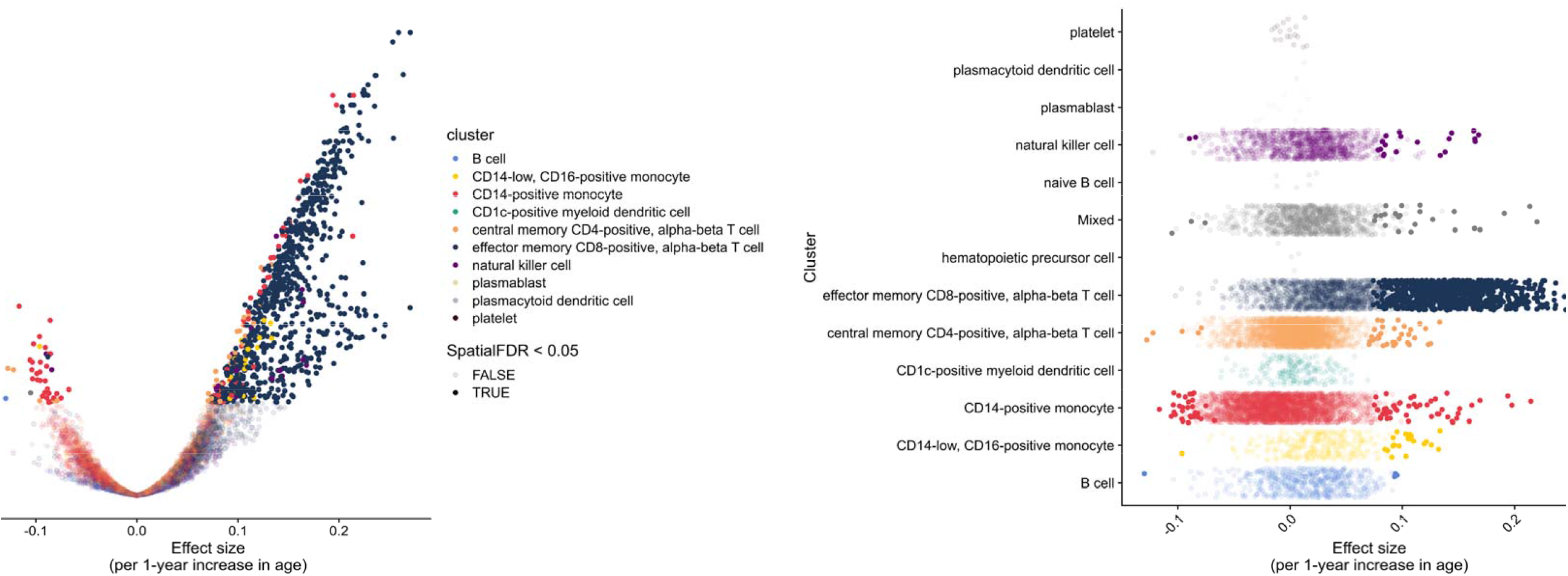
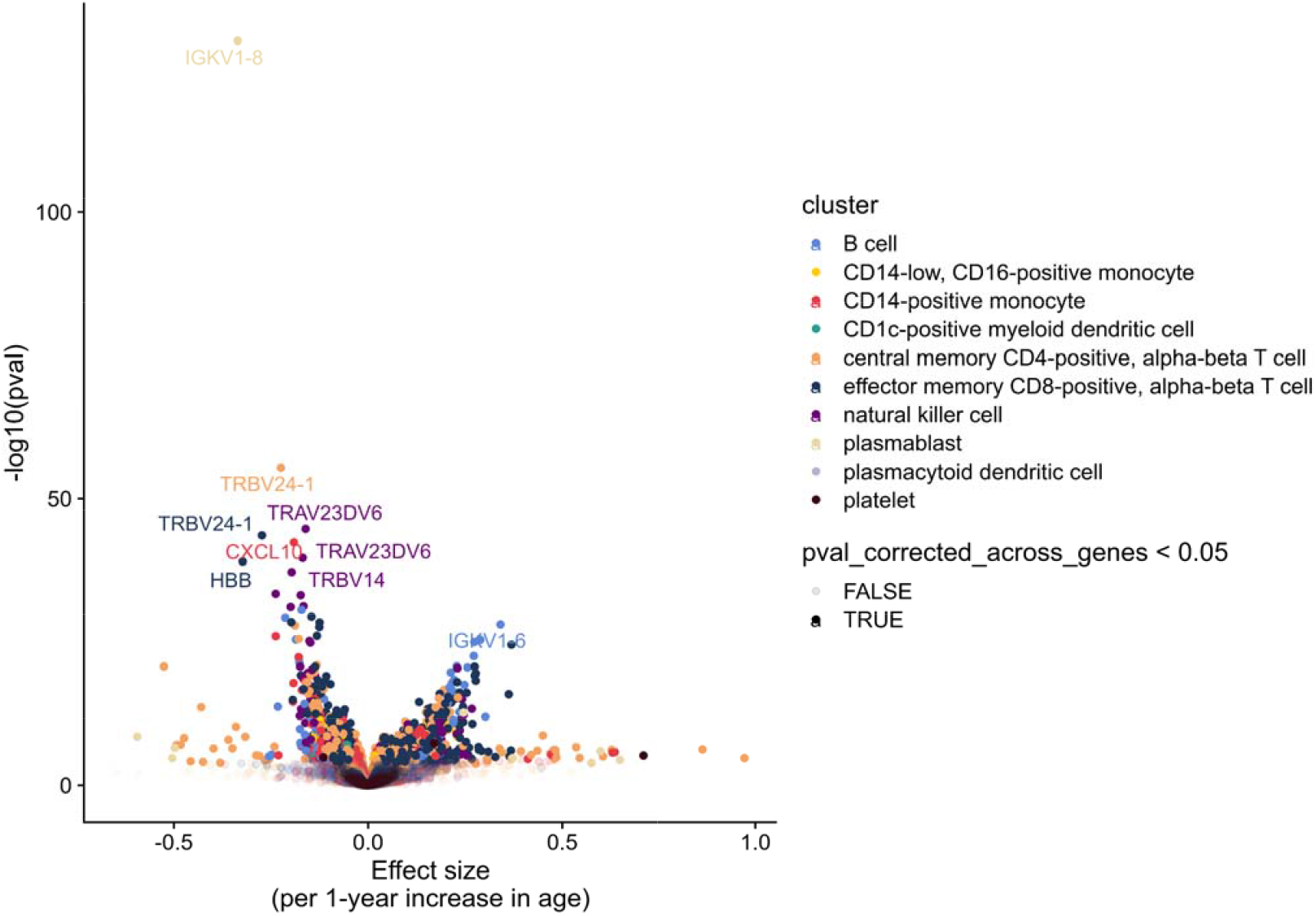
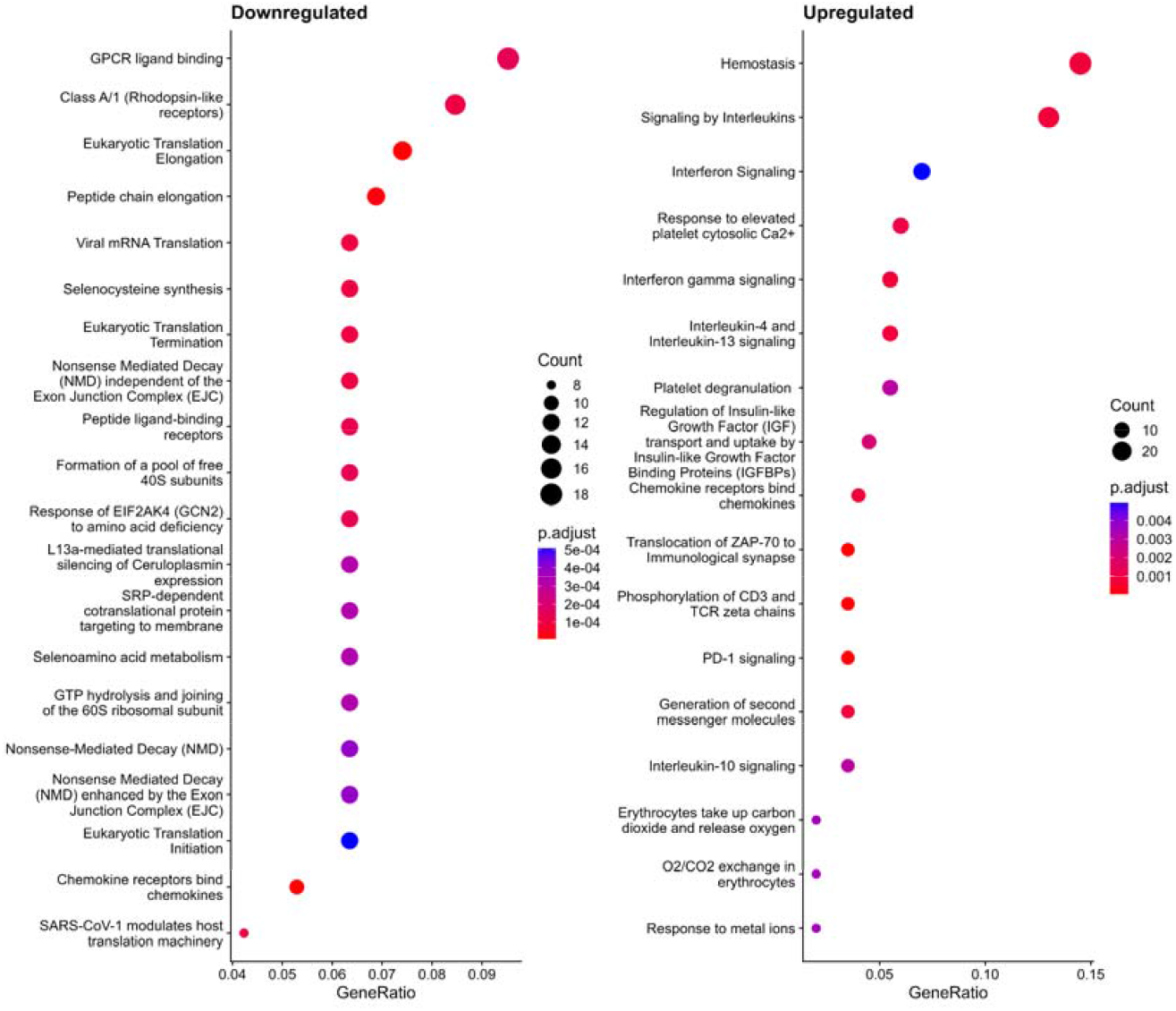
**(A)** Uniform manifold approximation and projection (UMAP), representing a total of 152,106 cells and 20 unique clusters. **(B)** *Left:* Volcano plot representing differential changes in neighborhoods across clusters per 1-year increase in recipient age. Effect sizes greater than 0 represent neighborhoods increasing in abundance with recipient age. Neighborhoods are colored by cluster and opaque dots represent differentially abundant neighborhoods (spatial FDR < 0.05). *Right:* Differential changes in neighborhoods represented in a beeswarm plot. Effect sizes greater that 0 represent neighborhoods increasing in abundance with recipient age, while opaque dots represent those neighborhoods that meet significance (spatial FDR < 0.05). **(C)** Volcano plot representing differentially expressed genes across neighborhoods. Genes are colored by cell type. An effect size greater than 0 represents genes with higher expression with increasing recipient age. Opaque dots represent those genes that met FDR < 0.05 within a tested neighborhood. A total of 878 unique differentially expressed genes were identified. **(D)** Reactome pathway enrichment for differentially expressed genes with increasing recipient age. Upregulated pathways include genes whose average logFC across neighborhoods was directionally positive with increasing recipient age, while downregulated pathways include genes whose average logFC across neighborhoods was directionally negative with increasing recipient age. With increasing recipient age, pathways involved in protein synthesis were downregulated while upregulated genes were enriched for pathways involving both immunosenescence (e.g., PD-1 signaling, autophagy, etc.) and chronic low-grade inflammation, consistent with dysregulated immunity.

Similarly, with increasing recipient age (adjusting for recipient sex and prednisone use), we found 878 unique differentially expressed (DE) genes in neighborhoods spanning 10 clusters (**Figure 3C, Supplemental Tables**). Beyond T cell receptor and immunoglobulin genes, a number of DE genes were associated with interferon signaling (*IFI27, IFI44/IFI44L, IFIT3, IFITM1, IFITM3* across *CD4+* T central memory cells, *CD8+* T effector memory cells, and *CD14+* monocytes) and inflammation (e.g., *CXCL8, CXCL9, CXCL10, CCL3, CCL4, CCL5*, etc.), with heterogeneity across neighborhoods. Of note, *CDKN1A*, which encodes p21 (key regulator of cell cycle progression and biomarker of cellular senescence)^15^, was significantly increased with increasing recipient age in *CD4+* and *CD8+* memory T cells, suggesting a potentially senescent phenotype with age. Pathway enrichment (**Figure 3D**) was performed by calculating the mean logFC for each significant gene (FDR < 0.05). Treating DE genes across all neighborhoods together (“pseudobulk”), downregulated genes were enriched for mechanisms of protein translation or ribosomal capacity, indicating an overall decreased protein synthesis capacity in aging cells. Simultaneously, upregulated genes were enriched for pathways involved both in immunosenescence (e.g., PD-1 signaling, autophagy, senescence-associated secretary phenotype) and chronic low-grade inflammation (e.g., interferon and interleukin signaling, platelet activation, etc.; **Supplemental Tables**).

## DISCUSSION

Our findings suggest a reduced risk of acute cardiac allograft rejection with increasing recipient age, with secondary findings of significant dysregulation of immune function with increasing age even in the context of immunosuppressive therapies after HT. We show evidence of increasing memory T cell compartments with recipient age, despite chronic illness and HT-associated immunosuppression, consistent with findings during aging in the non-transplant population^16^, and identified a number of DE genes enriched for immunosenescence and chronic low-grade inflammation pathways. While small clinical studies have previously observed reduced allograft rejection with increasing recipient age^17^, our work is a large, standardized, modern experience that integrates epidemiology and contemporary single-cell transcriptomic techniques. Key strengths of our study include a large clinical cohort spanning the modern era of HT for study, granular categorization of histological rejection grading (not typically available in national/international registries), and the use of single-cell approaches in HT recipients, an understudied area. Our single-cell data is reflective of “real world” cell-specific transcriptomic changes of those who develop cardiovascular disease in comparison to healthy aging atlases. For a field long overdue for personalization of care, these data suggest tangible strategies to mitigate HT-specific complications (e.g., rejection) while also reducing infections and malignancy driven by blanket immunosuppression for older individuals in HT.

The observation that older recipient age influences rejection risk with clear changes in immune cell phenotypes with aging calls for more personalized delivery of immunomodulation in an aging transplant population. These results underscore the importance of leveraging readily available clinical data to guide precision strategies in immunosuppression management, whether that may be rapid steroid withdrawal or targeting lower plasma levels of calcineurin inhibition. Such precision therapeutic approaches in the older transplant population can significantly improve both short- and long-term outcomes by balancing appropriate allograft protection while minimizing both immunological (e.g., infection, malignancy^13^) and non-immunological complications (cardiovascular-kidney-metabolic dysfunction, driven by immunosuppressive therapies^18^). Beyond chronological aging, approaches prioritizing epigenetic, transcriptomic, or proteomic aging clocks may further inform immunosuppressive strategies across the age span^19-22^.

Our study does have important limitations. Our sample size for single-cell RNA-sequencing comprises 40 individuals. Larger and more diverse data sets are essential to both validate our findings and uncover more subtle changes in gene signatures with aging.

Furthermore, studying the impact of aging on PBMCs in a multimorbid population (e.g., such as those warranting advanced cardiac therapies), rather than a healthy one as current atlases prioritize, and characterizing dynamic cell-specific transcriptomic changes that occur with immunosuppression initiation may more precisely inform therapeutic strategies.

In summary, our integrated clinical-translational study demonstrates that the risk of cardiac allograft rejection decreases with increasing recipient age at transplant, likely due in part to cell-specific transcriptional changes that occur with aging that reflect immunosenescence and dysregulated immunity. The convergence in our study of real-world clinical observations and single-cell profiling suggests an opportunity to better capture immunological function in older individuals in order to optimize outcomes after heart transplantation.

## METHODS

### Sex as a biologic variable

Our study examined male and female heart transplant recipients and similar findings are reported for both sexes.

### Study population and data collection

We conducted a retrospective analysis of all patients aged ≥18 years who underwent HT, inclusive of multi-organ transplant, between July 2013 and December 2023 at Vanderbilt University Medical Center (VUMC). Patients who were transplanted elsewhere prior to transfer of their care to VUMC were excluded due to lack of access to their endomyocardial biopsies. Similarly, recipients referred by the Veterans Affairs system who underwent HT at VUMC were excluded, as these patients receive all care following the index admission at the Veterans Administration hospital. This study was approved by the VUMC Institutional Review Board (IRB #200551).

Recipient data was extracted from the electronic health record and donor data was abstracted from the United Network for Organ Sharing (UNOS) database. Data collected included donor and recipient characteristics, donor cause of death, induction medications, and rejection type and grading. Following induction therapies peri-operatively, HT recipients at our center are maintained on an immunosuppressive regimen of calcineurin inhibitors, anti-proliferative agents, and steroids, which are typically weaned over the first 4-6 months following HT. Acute rejection was defined as clinically-relevant cellular or antibody-mediated rejection^11,12^: ACR grade 2R or 3R, pAMR1-i, pAMR2, and pAMR3.

### Statistical analysis (clinical-epidemiological)

Continuous variables are described as median (interquartile range; IQR) while categorical variables are described as N (%). Acute rejection episodes were restricted to within 1-year post-HT, as the overwhelming majority of endomyocardial biopsies occur within the first ∼6 months following HT and the risk of acute rejection drops significantly after the first year^10^. The mean and standard deviation (SD) for recipient age at transplant and donor age were calculated. The values were scaled and centered, so that the respective mean age was 0 and the SD was 1. Logistic regression was performed using the *rms* package (v 6.4-1) and lrm function. Covariates included donor age (in SD), recipient sex, and induction medications (with steroids alone as the reference). Effect sizes are represented per SD of age.

### Single-cell RNA-sequencing data set

We reprocessed and reanalyzed our recently-published single-cell RNA-sequencing (scRNA-seq) data set of peripheral blood mononuclear cells (PBMCs) obtained from HT recipients during routine clinical care (GSE271408)^14^. Briefly, samples were labeled with TotalSeq C hashtag antibodies for multiplexing and scRNA-seq data was generated using the 10X Genomics Single Cell 5’ v2 Reagent Kit. Sequencing was performed on the NovaSeq 6000 S4 flow cell targeting 50,000 reads per cell. The raw FASTQ files were aligned to the human genome (GRCh38-2024-A) and the RNA count matrix was generated using CellRanger (v9.0.1). Following importation into R (v4.4.2), ambient RNA was corrected using DecontX from Celda (v1.2.0). Cells were filtered by nFeature_RNA > 200 and < 3000. Multiplets were filtered out using HTODemux from Seurat (v5.0.2) and cells containing >5% mitochondrial genes were removed. A total of 152,106 cells passed quality control. Following normalization and batch correction, cell type annotation was performed by transferring labels from a publicly-available ScaleBio scRNA-seq data set available on CellXGene (https://cellxgene.cziscience.com/collections/4a9fd4d7-d870-4265-89a5-ad51ab811d89).

Differences in cell composition were conducted using Milo (v1.6.0)^23^, which constructs neighborhoods on the k-nearest neighbor (kNN) graph using a negative binomial generalized linear model and tests for differential abundance within each neighborhood. The kNN graph was constructed using *k* = 50, targeting ∼120 cells per neighborhood, and used 30 principal components (PCs). Recipient age was tested as a continuous variable and adjusted for recipient sex and prednisone use. Within each neighborhood, differential abundance was tested using the quasi-likelihood edgeR approach with multiple testing correction of nominal P-values performed using a weighted Benjamini-Hochberg method^24^. Effect sizes represent logFC per 1-year increase in recipient age. Neighborhoods were considered differentially abundant if they met a spatial false discovery rate (FDR)-adjusted P-value < 0.05.

Differential expression (DE) analysis was performed using miloDE (v0.0.0.9)^25^. Using this approach, transcriptionally-similar cells were assigned to neighborhoods using the 2^nd^ order kNN graph. We targeted a median of ∼1,280 cells per neighborhood, followed by DE analyses within each neighborhood using the quasi-likelihood edgeR approach. Using age as a continuous variable and adjusting for recipient sex and prednisone use, DE analyses were performed in 463 neighborhoods. Multiple testing correction of nominal P-values was performed using the Benjamini-Hochberg method for all genes tested both within each neighborhood and across all neighborhoods. DE genes were those that met an FDR-adjusted P-value < 0.05 within a neighborhood, with effect sizes representing logFC per 1-year increase in recipient age. Pathway enrichment analyses, with reactome terms prioritized, were performed using clusterProfiler (v4.6.2) using all tested genes as the background. Up- and down-regulated DE genes, regardless of neighborhood/cell type annotation, were defined by calculating the mean logFC across all neighborhoods. Multiple testing correction was conducted using the Benjamini-Hochberg method and reactome pathways were considered significantly enriched if meeting FDR-adjusted P-value < 0.05.

### Data and code availability

For the clinical-epidemiological analyses, data and code to confirm the findings of this study can be accessed upon reasonable request to the corresponding author (K.A.). For scRNA-seq analyses, code used for re-processing and analysis is available online at https://github.com/learning-MD/Aging_PBMC. Raw FASTQ files are deposited on the NCBI GEO (GSE271408).

## AUTHOR CONTRIBUTIONS

Conceptualization: K.A.; K.H.S., R.S., E.R.G.; Formal analysis: K.A., P.L., L.A.P., N.C., E.F.E., Q.S.W.; Writing and revision: K.A., K.H.S., R.S., H.K.S., J.E.F., E.R.G.

## REFERENCES

1. Lopez-Otin C, Blasco MA, Partridge L, Serrano M, Kroemer G. Hallmarks of aging: An expanding universe. Cell. 2023;186:243–278. doi: 10.1016/j.cell.2022.11.001

2. Gardner ID. The effect of aging on susceptibility to infection. Rev Infect Dis. 1980;2:801–810. doi: 10.1093/clinids/2.5.801

3. Li W, Zhang Z, Kumar S, Botey-Bataller J, Zoodsma M, Ehsani A, Zhan Q, Alaswad A, Zhou L, Grondman I, et al. Single-cell immune aging clocks reveal inter-individual heterogeneity during infection and vaccination. Nat Aging. 2025. doi: 10.1038/s43587-025-00819-z

4. Park MD, Le Berichel J, Hamon P, Wilk CM, Belabed M, Yatim N, Saffon A, Boumelha J, Falcomata C, Tepper A, et al. Hematopoietic aging promotes cancer by fueling IL-1⍰ - driven emergency myelopoiesis. Science. 2024;386:eadn0327. doi: 10.1126/science.adn0327

5. Erbe R, Wang Z, Wu S, Xiu J, Zaidi N, La J, Tuck D, Fillmore N, Giraldo NA, Topper M, et al. Evaluating the impact of age on immune checkpoint therapy biomarkers. Cell Rep. 2021;37:110033. doi: 10.1016/j.celrep.2021.110033

6. Nebhan CA, Cortellini A, Ma W, Ganta T, Song H, Ye F, Irlmeier R, Debnath N, Saeed A, Radford M, et al. Clinical Outcomes and Toxic Effects of Single-Agent Immune Checkpoint Inhibitors Among Patients Aged 80 Years or Older With Cancer: A Multicenter International Cohort Study. JAMA Oncol. 2021;7:1856–1861. doi: 10.1001/jamaoncol.2021.4960

7. Erbe R, Wang Z, Wu S, Xiu J, Zaidi N, La J, Tuck D, Fillmore N, Giraldo NA, Topper M, et al. Evaluating the impact of age on immune checkpoint therapy biomarkers. Cell Rep. 2021;36:109599. doi: 10.1016/j.celrep.2021.109599

8. Eochagain CM, Neuendorff NR, Gente K, Leipe J, Verhaert M, Sam C, de Glas N, Kadambi S, Canin B, Gomes F, et al. Management of immune checkpoint inhibitor-associated toxicities in older adults with cancer: recommendations from the International Society of Geriatric Oncology (SIOG). Lancet Oncol. 2025;26:e90–e102. doi: 10.1016/S1470-2045(24)00404-2

9. Colvin MM, Smith CA, Tullius SG, Goldstein DR. Aging and the immune response to organ transplantation. J Clin Invest. 2017;127:2523–2529. doi: 10.1172/JCI90601

10. Khush KK, Cherikh WS, Chambers DC, Harhay MO, Hayes D, Jr., Hsich E, Meiser B, Potena L, Robinson A, Rossano JW, et al. The International Thoracic Organ Transplant Registry of the International Society for Heart and Lung Transplantation: Thirty-sixth adult heart transplantation report - 2019; focus theme: Donor and recipient size match. J Heart Lung Transplant. 2019;38:1056–1066. doi: 10.1016/j.healun.2019.08.004

11. Stewart S, Winters GL, Fishbein MC, Tazelaar HD, Kobashigawa J, Abrams J, Andersen CB, Angelini A, Berry GJ, Burke MM, et al. Revision of the 1990 working formulation for the standardization of nomenclature in the diagnosis of heart rejection. J Heart Lung Transplant. 2005;24:1710–1720. doi: 10.1016/j.healun.2005.03.019

12. Colvin MM, Cook JL, Chang P, Francis G, Hsu DT, Kiernan MS, Kobashigawa JA, Lindenfeld J, Masri SC, Miller D, et al. Antibody-mediated rejection in cardiac transplantation: emerging knowledge in diagnosis and management: a scientific statement from the American Heart Association. Circulation. 2015;131:1608–1639. doi: 10.1161/CIR.0000000000000093

13. Youn JC, Stehlik J, Wilk AR, Cherikh W, Kim IC, Park GH, Lund LH, Eisen HJ, Kim DY, Lee SK, et al. Temporal Trends of De Novo Malignancy Development After Heart Transplantation. J Am Coll Cardiol. 2018;71:40–49. doi: 10.1016/j.jacc.2017.10.077

14. Amancherla K, Schlendorf KH, Chow N, Sheng Q, Freedman JE, Rathmell JC. Single-cell RNA-sequencing identifies unique cell-specific gene expression profiles in highgrade cardiac allograft vasculopathy. J Heart Lung Transplant. 2024. doi: 10.1016/j.healun.2024.11.017

15. Gasek NS, Yan P, Zhu J, Purushothaman KR, Kim T, Wang L, Wang B, Flynn WF, Sun M, Guo C, et al. Clearance of p21 highly expressing senescent cells accelerates cutaneous wound healing. Nat Aging. 2025;5:21–27. doi: 10.1038/s43587-024-00755-4

16. Terekhova M, Swain A, Bohacova P, Aladyeva E, Arthur L, Laha A, Mogilenko DA, Burdess S, Sukhov V, Kleverov D, et al. Single-cell atlas of healthy human blood unveils age-related loss of NKG2C(+)GZMB(-)CD8(+) memory T cells and accumulation of type 2 memory T cells. Immunity. 2023;56:2836–2854 e2839. doi: 10.1016/j.immuni.2023.10.013

17. Renlund DG, Gilbert EM, O’Connell JB, Gay WA, Jr., Jones KW, Burton NA, Doty DB, Karwande SV, Dewitt CW, Menlove RL, et al. Age-associated decline in cardiac allograft rejection. Am J Med. 1987;83:391–398. doi: 10.1016/0002-9343(87)90746-7

18. Huang S, Farber-Eger E, Tamaroff J, Chow N, Siddiqi HK, Brinkley DM, Menachem JN, Rali AS, Lindenfeld J, Ooi H, et al. Cardiovascular-kidney-metabolic disease burden in children and adults following heart transplantation. medRxiv. 2025.

19. Kabacik S, Lowe D, Fransen L, Leonard M, Ang SL, Whiteman C, Corsi S, Cohen H, Felton S, Bali R, et al. The relationship between epigenetic age and the hallmarks of aging in human cells. Nat Aging. 2022;2:484–493. doi: 10.1038/s43587-022-00220-0

20. Buckley MT, Sun ED, George BM, Liu L, Schaum N, Xu L, Reyes JM, Goodell MA, Weissman IL, Wyss-Coray T, et al. Cell-type-specific aging clocks to quantify aging and rejuvenation in neurogenic regions of the brain. Nat Aging. 2023;3:121–137. doi: 10.1038/s43587-022-00335-4

21. Argentieri MA, Xiao S, Bennett D, Winchester L, Nevado-Holgado AJ, Ghose U, Albukhari A, Yao P, Mazidi M, Lv J, et al. Proteomic aging clock predicts mortality and risk of common age-related diseases in diverse populations. Nat Med. 2024;30:2450–2460. doi: 10.1038/s41591-024-03164-7

22. Oh HS, Rutledge J, Nachun D, Palovics R, Abiose O, Moran-Losada P, Channappa D, Urey DY, Kim K, Sung YJ, et al. Organ aging signatures in the plasma proteome track health and disease. Nature. 2023;624:164–172. doi: 10.1038/s41586-023-06802-1

23. Dann E, Henderson NC, Teichmann SA, Morgan MD, Marioni JC. Differential abundance testing on single-cell data using k-nearest neighbor graphs. Nat Biotechnol. 2022;40:245–253. doi: 10.1038/s41587-021-01033-z

24. Lun ATL, Richard AC, Marioni JC. Testing for differential abundance in mass cytometry data. Nat Methods. 2017;14:707–709. doi: 10.1038/nmeth.4295

25. Missarova A, Dann E, Rosen L, Satija R, Marioni J. Leveraging neighborhood representations of single-cell data to achieve sensitive DE testing with miloDE. Genome Biol. 2024;25:189. doi: 10.1186/s13059-024-03334-3

